# Accounting for imperfect detection in species with sessile life cycle stages: a case study of bumble bee colonies

**DOI:** 10.1101/518407

**Authors:** David T. Iles, Genevieve Pugesek, Natalie Z. Kerr, Nicholas N. Dorian, Elizabeth E. Crone

## Abstract

1. For bumble bees, colonies (not individual workers) are the functional unit of the population. Estimates of colony density are thus critical for understanding population distribution and trends of this important pollinator group. Yet, surveys of bumble bee colonies and other taxa with sessile life cycle states rarely account for imperfect detection.
2. Here we demonstrate the use of mark-recapture methods to estimate the density of bumble bee colonies across the landscape using standardized survey protocols.
3. We found that the probability of detecting colonies in standardized surveys varied considerably across space, through time, and among colonies.
4. Using simulations, we also show that imperfect detection can obscure true variation in density among plots, or generate spurious variation in counts even when all plots have the same density. In both cases, we show that mark-recapture can be used to generate unbiased estimates of density, with relatively low search effort compared to conventional survey methods for bumble bee colonies.
5. Our study illustrates the advantages of mark-recapture for optimizing survey protocols for species with cryptic and sessile life cycle stages, which will be a valuable tool in ongoing studies of pollinator nesting ecology.

## Introduction

Measuring the size of natural populations is a primary goal of conservation monitoring and is often a prerequisite for management action or wildlife policy decisions (Williams et al. 2002). Yet, estimating abundance and the factors that influence it can be challenging when organisms are difficult to detect. Accordingly, study designs and analyses that account for imperfect detection have a rich tradition in wildlife research (White and Burnham 1999; Williams et al. 2002; Kéry and Schaub 2012). Failure to account for imperfect detection of cryptic organisms can introduce both uncertainty and bias into population estimates, potentially leading to erroneous conclusions about population status and habitat requirements (Gu and Swihart 2004; Kéry and Schmidt 2008).

In contrast to studies of mobile vertebrates, imperfect detection is rarely accounted for in population estimates for both invertebrates and taxa that are sessile or have a sessile life-cycle stage, including plants and ground-nesting animals (Kellner and Swihart 2014; Berberich et al. 2016). Kellner and Swihart (2014) found that only 9.0% of invertebrate and 1.4% of plant population studies accounted for imperfect detection, compared to the mean of 23% across all taxa. This discrepancy potentially stems from the incorrect assumption that detection probability is inherently high in less mobile organisms. Yet, this assumption may be false if sessile organisms are rare, inconspicuous, or logistically difficult to survey. For example, in a long-term study of a 4.5 ha plot in Kansas, Slade et al. (2003) estimated that fewer than 4% of existing Mead’s milkweed plants were discovered in years without spring fires, and only 18% were discovered in years with spring burning. In a study of Baltimore Checkerspot butterfly demography, Brown et al. (2017) estimated the density of caterpillar aggregations on the landscape using mark-resight surveys. The detection probability ranged from 0.32 to 0.56 on native *Chelone glabra* host plants, and from 0.17 to 0.25 on exotic *Plantago lanceolata* plants (L.M. Browne and E.E. Crone, pers. comm, from analyses in Brown et al. 2017). Surprisingly, Berberich et al. (2016) found that observers conducting one hour surveys for red wood ant nests in 60m × 60m plots failed to detect up to 40% of ant nests larger than 50cm. Thus, even for studies of sessile organisms, detection can be highly imperfect and systematically biased.

Bumble bees are an important group of pollinators with a sessile life-cycle stage. After a queen bumble bee has established a colony and produced her first cohort or two of workers, she remains in the nest until the colony expires (Goulson 2010). Since colonies are the functional unit of the population for social insects such as bumble bees, studies of nesting habitat are particularly valuable for conservation planning. Yet, colonies are difficult to find, so relatively few studies have investigated the correlates of nest density compared to the numerous studies of habitat preference by foraging workers (but see examples in Harder 1986; Osborne et al. 2008; Waters et al. 2010; Lye et al. 2012; O’Connor et al. 2012, 2017). This disparity is problematic because workers are highly mobile, e.g., workers often forage up to 1 km from their colony (Greenleaf et al. 2007) and may be attracted to areas rich in floral resources. This decoupling of foraging and nesting sites potentially obscures the landscape drivers of population performance (Heard et al. 2007). Indeed, Herrmann et al. (2007) reported that bumble bee colony densities were uncorrelated with local worker densities across an agricultural landscape in Germany.

A variety of methods have been used to estimate the density of bumble bee nests across landscapes. These methods have included “free searches” in which observers haphazardly search particular habitat types for nests (Harder 1986; O’Connor et al. 2012; Rao and Skyrm 2013; O’Connor et al. 2017), canine-assisted searches (Waters et al. 2010; O’Connor et al. 2012), and concentrated stationary observation of small plots by individual researchers or distributed citizen science networks (Osborne et al. 2008; Lye et al. 2012). As an alternative to ground-based searches, molecular (i.e., genetic) analysis of foraging workers has also been used to infer nest density at large spatial scales (Darvill et al. 2004; Goulson et al. 2010). Other studies have inferred the relative density of nests across space based on the prevalence of spring prospecting behavior by newly emerged queens (Svensson et al. 2000; Kells and Goulson 2003; O’Connor et al. 2017). Combined, these studies have reported considerable variation in nest densities (range: 0.1 to 50.1 nests· ha^−1^; Table 2 in Appendix 1), potentially owing to ecologically relevant differences across species, habitat types, and landscape configurations. However, O’Connor et al. (2012) also reported a 20-fold difference in the number of nests detected between fixed and free searches, indicating an extreme degree of variation in nest detection among survey strategies. Detection probability also often varies through time, across space, and between individuals (Anderson 2001; Kéry and Schaub 2012). Therefore, unaccounted differences in detection probability within and among studies could contribute to observed variation in nest density, limiting the ability to generalize across studies and resolve the true environmental correlates of bumble bee population abundance.

Here, our objective was to generate unbiased estimates of bumble bee nest density using mark-recapture methods while simultaneously examining the factors that influence imperfect detection of nests. We first discuss the general application of closed-population modelling approaches to estimate the abundance of cryptic sessile organisms. We then review a classic catalogue of model structures that can be used to correct for systematic bias in detection probability, along with their specific relevance to our study of bumble bee nest density. We fit these models to empirical data and examine the consequences of imperfect nest detection in a field setting. Finally, we conduct a simulation to illustrate the (spurious) variability induced into count data by imperfect detection, and to demonstrate that this approach can be usefully applied in cases where a large number of sites are only visited twice. Our study emphasizes the utility of mark-recapture approaches for examining the ecological correlates of nest density for social insects, and outlines a strategy for surveying bumble bee colonies with imperfect detection.

### Mark-recapture for estimating abundance with imperfect detection

Closed population models can be used to estimate abundance while accounting for imperfect detection when the processes of birth, mortality, and movement do not alter the number of individuals in plots over the course of a study (i.e., when the population is “demographically closed” across the sampling period). For sessile organisms, this assumption is satisfied when birth and death processes are unlikely to occur across the sampling period. Individuals can also be censored if they are known to have died during the sampling period.

Otis et al. (1978) outlined a classic catalogue of model structures that can be used to examine drivers of variation in detection probability for closed populations. We adopt this framework for our analysis of bumble bee nests to demonstrate how these models can be applied to studies of sessile organisms and to link our specific study system to a well-defined body of mark-recapture research. Mathematical descriptions of each model are presented in Table 1.

**Table 1.**
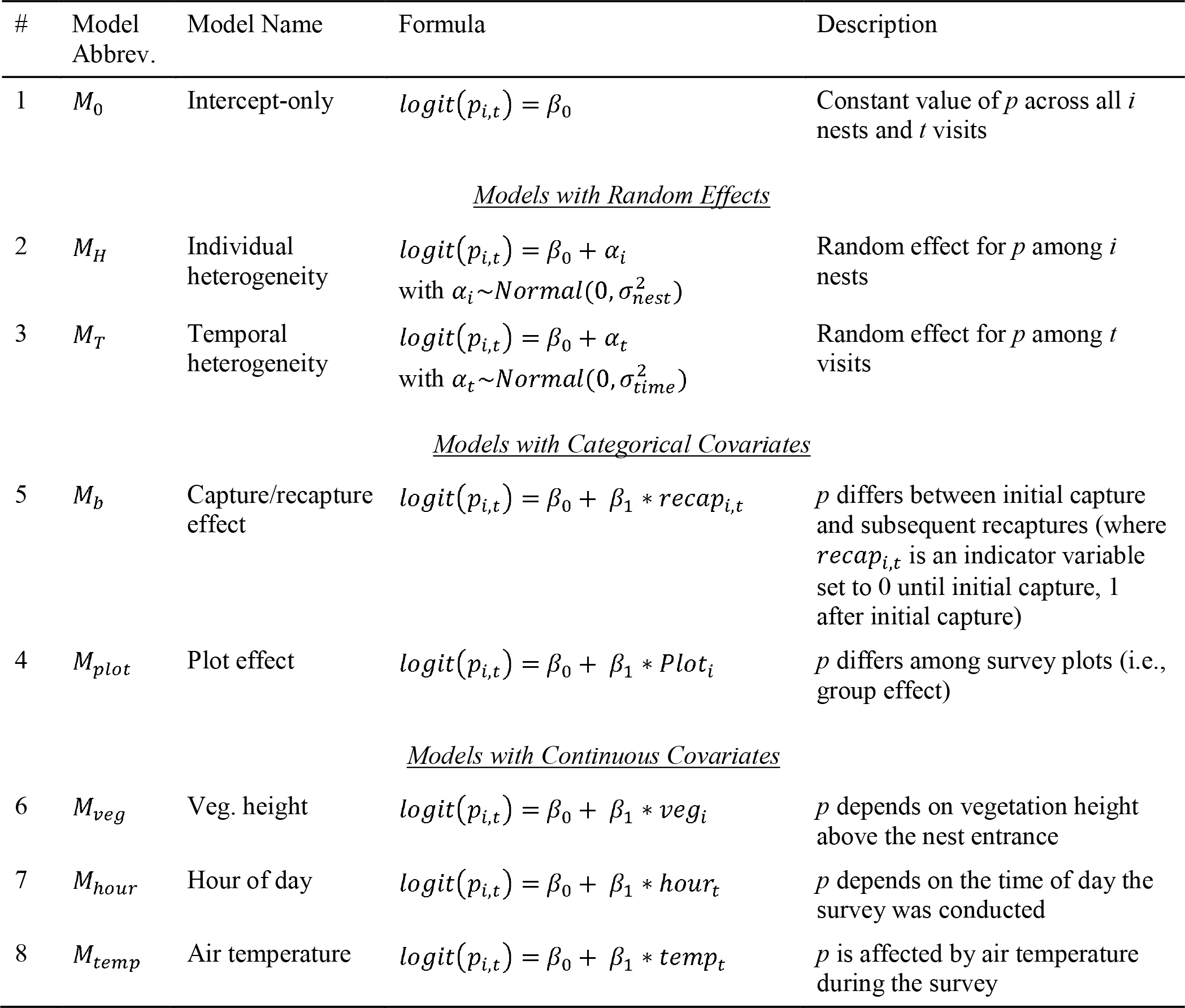
Classic closed population model structures used in our study to examine variation in the probability of detecting bumble bee nests. *p*_*i*,*t*_ refers to the detection probability of nest *i* on survey *t*, and *β*_*i*_ refer to various fitted model coefficients

| # | Model Abbrev. | Model Name | Formula | Description |
| --- | --- | --- | --- | --- |
| 1 | $M_0$ | Intercept-only | $\text{logit}(p_{i,t}) = \beta_0$ | Constant value of $p$ across all $i$ nests and $t$ visits |
| <u>Models with Random Effects</u> |  |  |  |  |
| 2 | $M_H$ | Individual heterogeneity | $\text{logit}(p_{i,t}) = \beta_0 + \alpha_i$<br>with $\alpha_i \sim \text{Normal}(0, \sigma_{nest}^2)$ | Random effect for $p$ among $i$ nests |
| 3 | $M_T$ | Temporal heterogeneity | $\text{logit}(p_{i,t}) = \beta_0 + \alpha_t$<br>with $\alpha_t \sim \text{Normal}(0, \sigma_{time}^2)$ | Random effect for $p$ among $t$ visits |
| <u>Models with Categorical Covariates</u> |  |  |  |  |
| 5 | $M_b$ | Capture/recapture effect | $\text{logit}(p_{i,t}) = \beta_0 + \beta_1 * \text{recap}_{i,t}$ | $p$ differs between initial capture and subsequent recaptures (where $\text{recap}_{i,t}$ is an indicator variable set to 0 until initial capture, 1 after initial capture) |
| 4 | $M_{plot}$ | Plot effect | $\text{logit}(p_{i,t}) = \beta_0 + \beta_1 * \text{Plot}_i$ | $p$ differs among survey plots (i.e., group effect) |
| <u>Models with Continuous Covariates</u> |  |  |  |  |
| 6 | $M_{veg}$ | Veg. height | $\text{logit}(p_{i,t}) = \beta_0 + \beta_1 * \text{veg}_i$ | $p$ depends on vegetation height above the nest entrance |
| 7 | $M_{hour}$ | Hour of day | $\text{logit}(p_{i,t}) = \beta_0 + \beta_1 * \text{hour}_t$ | $p$ depends on the time of day the survey was conducted |
| 8 | $M_{temp}$ | Air temperature | $\text{logit}(p_{i,t}) = \beta_0 + \beta_1 * \text{temp}_t$ | $p$ is affected by air temperature during the survey |

The simplest model, *M*_0_, estimates a single detection probability that is common across all nests and sampling occasions. In reality, individuals may differ in their probability of being detected, a phenomenon known as individual heterogeneity. Consequently, the model *M*_*H*_ estimates both a mean and variance in detection probabilities across nests (i.e., individual random effects). In our study of bumble bee nests, this could be due to unmeasured differences in the worker activity level or the location of nests that increase the probability particular nests will be detected. Similarly, the model *M*_*T*_ estimates a mean detection probability across nests and a variance in detection associated with visits (i.e., a temporal random effect). For bumble bees, this is could be due to temporal variation in colony size, ambient temperature (which potentially affects activity level), or seasonal changes in vegetation within plots that alter the probability nests will be detected on each visit. The model *M*_*b*_ described by Otis et al. (1978) accounts for a discrete behavioral change in organisms that affects their individual detection in subsequent marking occasions, commonly referred to as “trap-shyness” or “trap-happiness”. In studies of sessile organisms, rather than behavioral changes of the study organisms themselves, this response can plausibly occur if the vegetation surrounding focal organisms becomes trampled by researchers or if researchers remember the location of individuals (in this case, nests) they have previously located. Either of these scenarios would result in different detection probabilities for initial and subsequent capture events in studies of sessile organisms.

Although not explicitly described by Otis et al. (1978), fixed effects of explicit covariates can also be incorporated to examine the drivers of variation in detection probability (Kéry and Schaub 2012). These can include age or size of the organism, habitat covariates, or explicit temporal covariates (e.g., to examine temporal trends in detection). In our study, four of the eight models included explicit covariates affecting detection of bumble bee nests. Towards this goal, we constructed a model that included different detection probabilities for each survey plot (*M*_*plot*_; see *Study Site* in *Method*s section below), and three models that included effects of vegetation height above the nest (*M*_*veg*_), hour of the day at which plots were surveyed (*M*_*hour*_), and ambient air temperature during the survey (*M*_*temp*_). Our objective was to demonstrate how these effects could be incorporated to generate deeper insights into the processes influencing nest abundance surveys, rather than to exhaustively examine the diverse suite of (potentially interacting) factors that influence nest detection.

## Methods

### Study Site

We conducted searches for bumble bee nests in three survey plots located at Appleton Farms (42.65°N, 70.86°W) in Ipswich, Massachusetts. Each plot was approximately 3000 m^2^ in area and not used for agriculture, although each plot is mowed annually to prevent succession. Two plots were adjacent to one another, while the third plot was located approximately 1,000 meters away. Primary vegetation cover in each plot consisted of a variety of grasses, sedges (*Carex* spp.), perennial forbs (e.g., *Plantago lanceolata*, *Linaria vulgaris*, *Lotus corniculatus*, *Solidago* spp., *Asclepias* spp.), and ericaceous shrubs (e.g., *Vaccinium angustifolium*, *Rubus* spp.). Each plot was bordered by a hedgerow of trees or forest, and the surrounding landscape was mixed agriculture (pasture and hay fields) and natural areas (forest and wet meadows).

### Data Collection

Each plot was visited repeatedly in July and August of 2017, after bumble bee colonies had produced several cohorts of workers, between 6:30 am and 7:30 pm, though the majority of nest surveys took place in the morning. Searches were conducted independently by eight different investigators, several of whom surveyed each plot multiple times. Each plot was surveyed for two hours between 12-16 times, for a total effort of 24-32 survey hours per plot. During each survey, searchers moved slowly through the plot looking for bumble bee activity that might indicate the presence of nest (i.e., workers quickly descending to the ground, slowly ascending from the ground, or conducting circular navigation flight behavior). Upon locating a bumble bee nest, searchers placed an inconspicuous, numbered identifier next to the nest entrance and recorded the nest location, identity, and whether the nest had been previously located either by themselves or by other searchers. For each located nest, we recorded the height of the tallest vegetation immediately above the nest entrance, which could affect searcher ability to detect bumble bee movement near nest entrances. We extracted hourly ambient air temperature measurements during each survey from the nearest weather station (approx. 9.5 km away); ambient air temperature could affect bumble bee metabolism and nest activity, which could influence our ability to detect nests. Finally, we also recorded the time of day of each survey, as bumble bee activity varies throughout the day (Kwon and Saeed 2003).

### Statistical Methods

The foundation of mark-recapture approaches is the encounter histories of individuals (in this case bumble bee colonies) that are generated from repeated surveys of plots. Closed population models assume the abundance of individuals within each plot does not change across sampling periods; thus, variation in the number of individuals detected across repeated surveys is caused entirely by observation error. The goal of closed population models is to estimate detection probability (*p*) of individuals along with the spatial, temporal, and/or individual-level factors that influence it. Once the factors that influence detection probability are estimated, the observed count of individuals can be corrected to generate an unbiased estimate of the true number of individuals present at a site.

Successful detection of the individual *i* during each survey *t* occurs with some probability *p*_*i*,*t*_ and the probability of not detecting the individual is 1 − *p*_*i*,*t*_. An encounter history of ‘0110’ implies the individual was detected on the second and third survey of the plot and not detected on the first and fourth survey. The entire encounter history ‘0110’ for individual *i* therefore occurs with probability (1 − *p*_*i*,1_) × *p*_*i*,2_ × *p*_*i*,3_ × (1 − *p*_*i*,4_). The actual encounter history (i.e., the observed sequence of 0’s and 1’s) is assumed to arise from a series of Bernoulli trials. Additionally, a link function can be used to model the influence of covariates on detection probability, analogous to a logistic regression.

We fit the series of closed population models described in Table 1 using the empirical nest encounter histories generated by our repeated surveys of plots. We fit models using Bayesian methods, outlined by Kéry and Schaub (2012, ch 6); though we note that such models can also be fit in a frequentist framework using maximum likelihood approaches. Bayesian analysis allows for random effects to be easily incorporated, for nests to be right-censored part way through the study (e.g., if a nest was known to have failed, which violates an assumption of closed population models), and for uncertainty in parameter estimates to be easily propagated to the model output. The Bayesian implementation of closed population models uses an additional technique called “data augmentation” to estimate the true number of individuals in a plot, based on the number of nests actually detected and their estimated detection probabilities (see Kéry and Schaub 2012, ch. 6; Royle and Dorazio 2008, section 5.6 for further discussion of this technique; also see our implementation of this approach in Supplementary Material 1).

### Simulation Study

The methods described above demonstrate the application of a mark-recapture approach to the study of sessile insect life cycle stages when detection is highly imperfect. However, to further illustrate the value of this approach, we conducted a simulation to illustrate a second (and converse) problem of failing to account for imperfect detection: imperfect detection generates spurious sampling variability in counts, even when no variation exists. This simulation was also used to examine whether the models we used would converge with only two visits (the minimum number of visits required for mark-recapture) if a larger number of plots had been surveyed. We simulated 40 hypothetical plots, each containing exactly 5 bumble bee nests. We used a fixed detection probability of 0.30 for all nests (approximately equal to the mean detection probability we estimated; see Results section and Fig. 1a). We then simulated the number of nests observed on a single visit to each plot by drawing from a binomial distribution with 5 trials for each plot (i.e., one for each nest). We used this simulation to evaluate 1) the degree of spurious variability introduced into counts simply by imperfect detection, and 2) whether the closed population mark-recapture approach could correct for this spurious variability with only two visits to each plot.

**Fig. 1.**
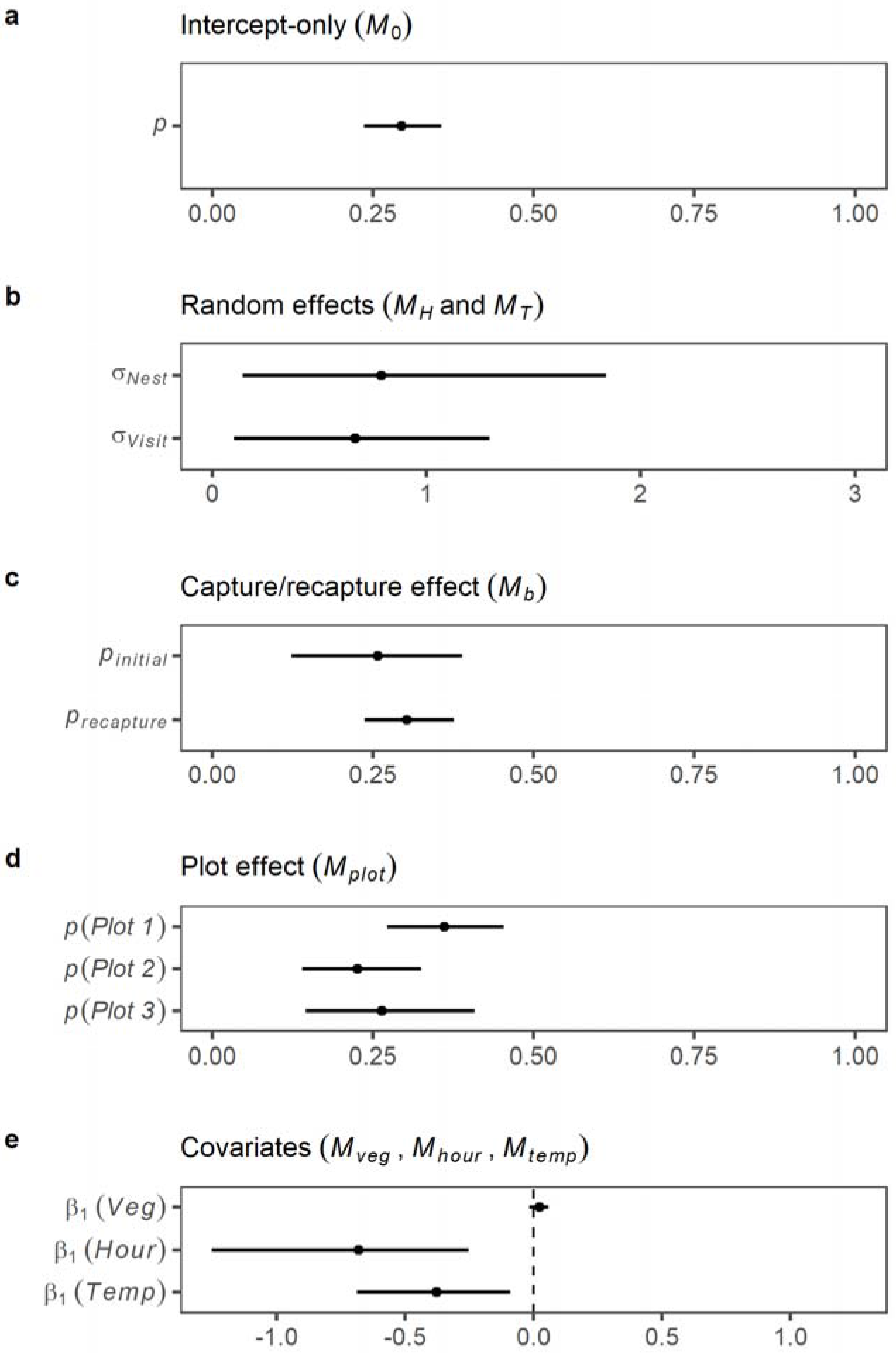
Effects on bumble bee nest detection probability from closed population models (n = 18 nests). Points represent median estimate of effect from Bayesian posterior distributions; lines represent 95% credible intervals

To facilitate the straightforward application of this analytical toolkit to other studies of sessile insect life cycle stages, we include full encounter history data, covariate data, and commented R code for Bayesian analysis and simulations as Supplementary Materials 1-9. Data management and simulations were conducted in R version 3.4.4 and Bayesian analysis was conducted in JAGS using the *jagsUI* library in R (Kellner 2018).

## Results

We located 18 bumble bee nests across the three survey plots (10, 5, and 3 nests in each plot, respectively). All nests were constructed by *Bombus impatiens*, except for one that was constructed by *B. bimaculatus*. We used all nests for subsequent analysis. The number of nests located on single visits to each of the three plots ranged from 0 to 6, 0 to 3, and 0 to 3 for each plot, respectively. The three plots were searched 11, 16, and 14 times by at least 6 different observers.

### Model results – patterns in nest detection

The mean detection probability of nests based on the intercept-only model (*M*_0_) was 0.30 (95% credible interval [CRI] = 0.24 − 0.36; Fig. 1a). A model that included heterogeneity in *p* across nests (*M*_*H*_) suggested that nests differed in their individual detection probabilities, with a median posterior estimate for *p* of 0.26 (95% CRI = 0.11 to 0.37) and standard deviation (on the logit scale) of 0.79 (95% CRI = 0.14 to 1.84; Fig. 1b). Similarly, a model that included heterogeneity in *p* across visits (*M*_*T*_) indicated that detection probability varied through time with a median *p* of 0.27 (95% CRI = 0.20 to 0.38) and standard deviation (on the logit scale) of 0.67 (95% CRI = 0.10 to 1.29; Fig. 1b). We note that there was substantial uncertainty associated with estimates of both individual and temporal random effects, as is common for random effect models fit to relatively sparse data (Kéry and Schaub 2012).

There was weak evidence for different detection probabilities between the first and subsequent capture occasions (Fig. 1c). Thus, nests were not more likely to be detected after their initial discovery.

A model including different detection probabilities for nests within each survey plot (*M*_*plot*_) indicated nest detectability varied systematically across plots (Fig. 1d). Under this model, median estimates of detection probabilities in each plot were 0.36 (95% CRI = 0.27 to 0.45), 0.23 (95% CRI = 0.14 to 0.33), and 0.26 (95% CRI = 0.15 to 0.41). The probability that detection probability was greater for nests in plot 1 than plot 2 was 0.98 (calculated directly from posterior probability distributions).

We then constructed a series of models to examine effects of specific covariates on *p*. Height of vegetation above the nest entrance did not have a strong effect on *p* (standardized effect of vegetation height from *M*_*veg*_ = 0.02; 95% CRI = −0.02 to 0.06; Fig. 1e). However, *p* declined strongly throughout the day during sampling times (standardized effect of hour from *M*_*hour*_ = −0.68; 95% CRI = −1.25 to −0.25; Fig. 1e) and was negatively correlated with ambient air temperature during the survey (standardized effect of air temperature from *M*_*temp*_ = −0.38; 95% CRI = −0.69 to −0.09; Fig. 1e).

### Model results – nest abundance

Across all eight models, median estimates of nest abundance were in close agreement. All models estimated approximately 10, 5, and 3 nests in each of the three plots, respectively (Fig. 2). The corresponding median estimate of nest density in each plot was therefore 33.3, 16.7, and 10 nests·ha^−1^. Consequently, on single surveys of each plot, we located approximately 0-60% of the nests in plots 1 and 2, and 0-100% of the nests in plot 3.

**Fig. 2.**
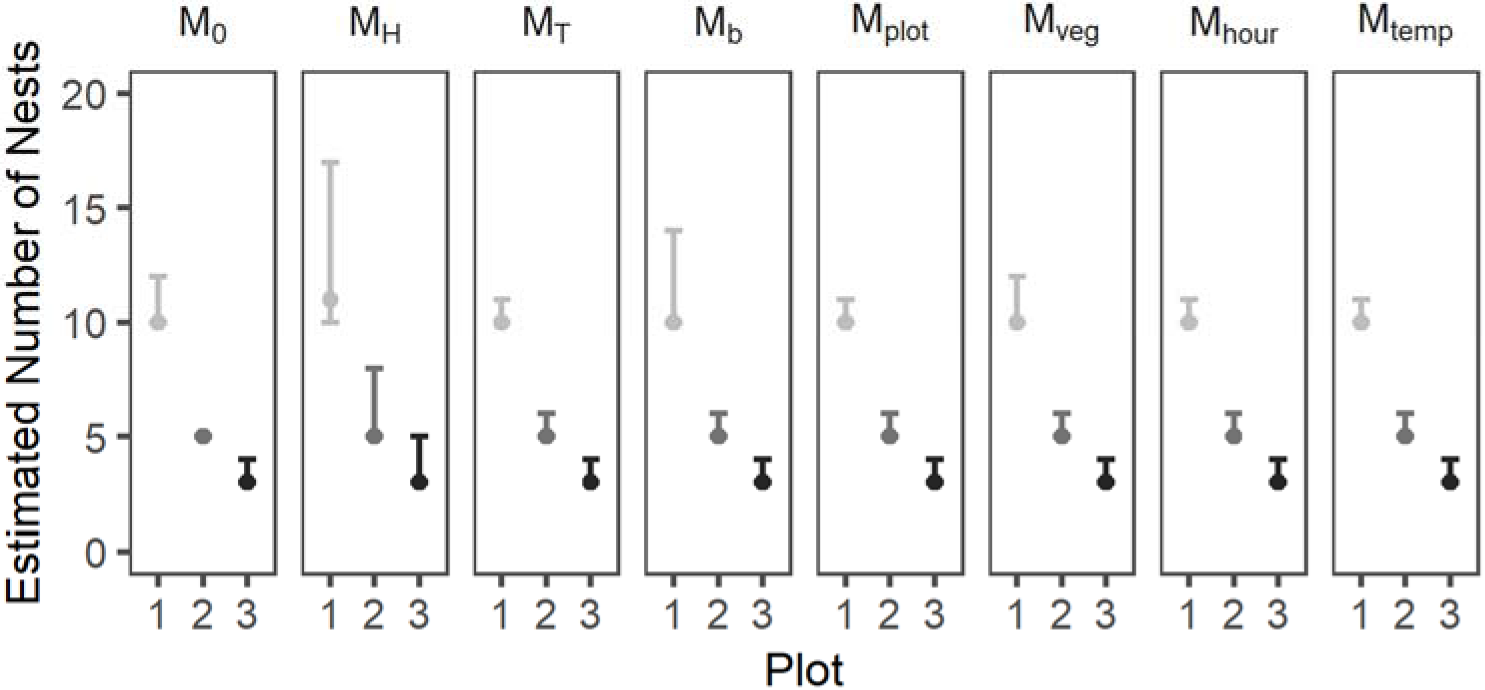
Estimated abundance of nests in each of the three survey plots (with associated 95% Bayesian credible intervals) based on each of the eight closed population models and all survey data

However, even with our large number of repeated searches (11-16 per survey plot), there was a high probability that undiscovered nests remained in each plot at the end of our study (Fig. 2; note range of credible intervals). For example, while the median estimate of total nest abundance from model *M*_0_ was 18 (equal to the number of nests we located), the 95% credible interval was 18 to 20, and the probability that the true abundance was greater than 18 was 0.25. Notably, the credible intervals for estimates from model *M*_*H*_ were wide relative to other models. This reflects two important features of individual heterogeneity: 1) a fraction of nests have extremely low detection probabilities and it is difficult to estimate how many remained undetected, and 2) the existing amount of heterogeneity is difficult to estimate, especially with low sample sizes (see Fig. 1b).

Given that our counts of bumble bee nests were subject to substantial observation error, we performed an additional analysis to illustrate how imperfect detection can obscure large differences in the density of colonies among plots. We randomly selected nest counts from single surveys to each plot, calculated the resulting rank order of nest densities, and compared them to our estimates from mark-recapture models. We repeated this process 1000 times. This analysis revealed that based on single visits to each field, the incorrect rank-order of density between plots arises 70% of the time despite large differences in the relative density of nests in each plot. Strikingly, the incorrect rank order between plots 1 and 3 arises 17% of the time, despite a three-fold difference in estimated nest density between these two plots (33.3 vs 10 nests·ha^−1^). Next, to quantify the effort needed to reliably estimate differences in nest density between plots, we sequentially re-fit model M_0_ for different numbers of visits. With our small number of plots and so few nests initially detected, the model would not converge with only 2 visits to each plot. This also occurred when models were fit with maximum likelihood in program MARK instead of using Bayesian methods. The model converged with 3 visits to each plot, but uncertainty associated with abundance estimates was extremely large (Fig. 3). After 5 visits to each plot, clear differences in abundance between plots 1 and 3 were apparent. As expected, uncertainty in estimates continued to decline as the number of surveys increased.

**Fig. 3.**
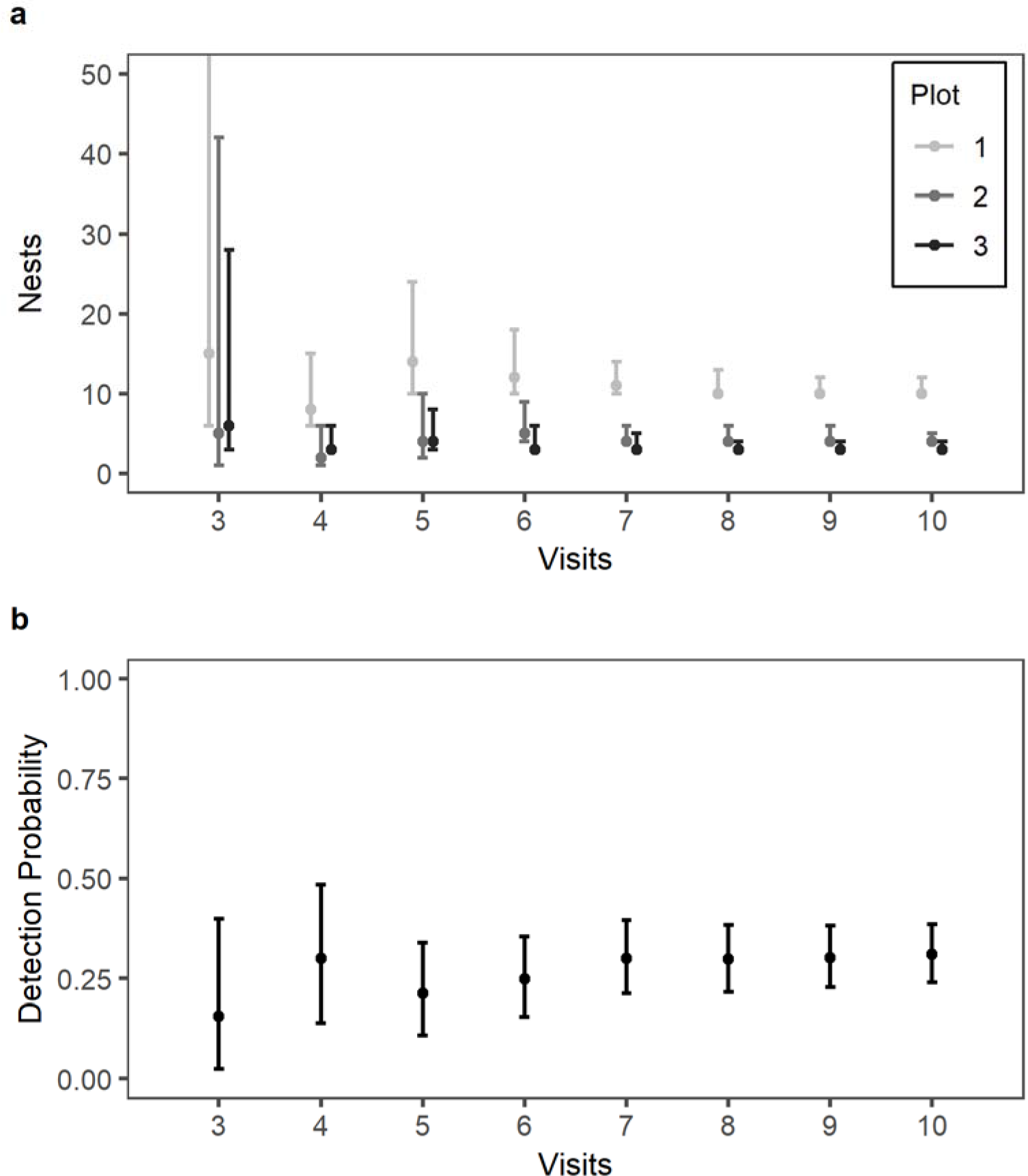
Estimated nests in each plot and associated mean detection probability from closed population models, based on model M_0_ after different numbers of visits to each plot. Points represent median estimates from Bayesian posterior distribution; lines denote associated 95% credible intervals

### Simulation results – correcting spurious variation in nest abundance

Our final simulation illustrates the sampling variability induced into count data even when detection rates are fixed at 0.3. We then used this simulation to examine whether this spurious source of variability could be corrected by visiting each plot only twice if a larger number of plots were visited. Despite the true presence of exactly 5 nests in each of the 40 simulated plots (Fig. 4; solid black line), the number of nests detected per plot ranged from 0 to 4 with a mean of 1.51 and standard deviation across plots of 0.95 (Fig. 4, dashed line and dots). Thus, based on a single visit to each plot, there is considerable (but spurious) variation in nest abundance across the 40 plots. We then simulated a second visit to each plot, fit mark-recapture model *M*_0_ to the resulting encounter histories, and estimated the number of nests in each plot while accounting for imperfect detection. The model successfully converged with only 2 visits to each plot and the credible intervals (Fig. 4; gray ribbons) overlapped the true abundance in every plot. The model *M*_0_ therefore correctly indicates weak evidence for variation in abundance among the 40 simulated plots, despite the high degree of variation in nest counts from a single survey. Thus, for larger sample sizes of plots, two to three visits to each may be sufficient in estimating nest densities.

**Fig. 4.**
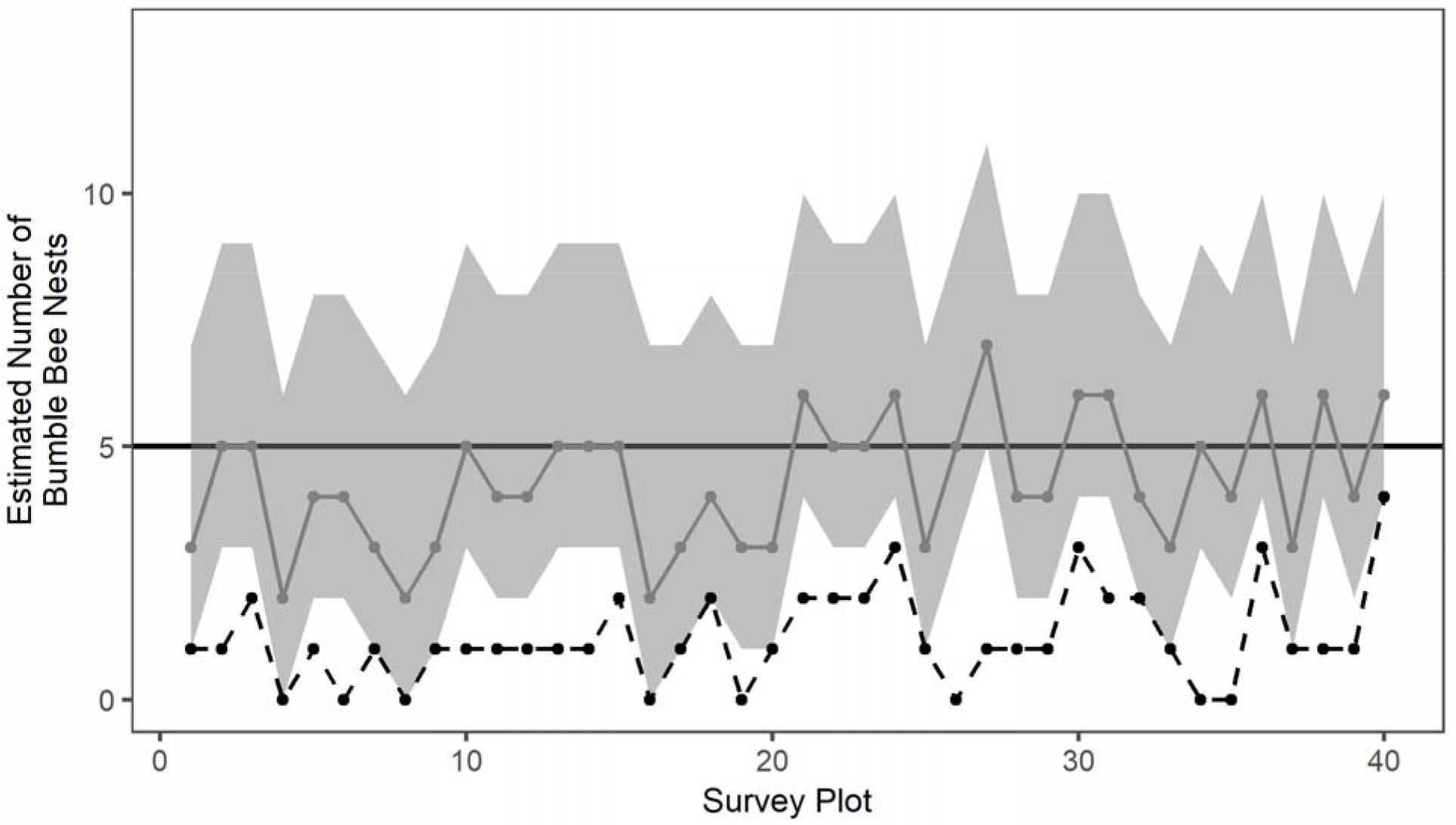
Results of a simulation with 40 survey plots, each with exactly 5 nests, and each nest with a 0.3 probability of detection on a single visit. Black dashed lines and black dots represent counts on a single visit to each plot. Gray lines and dots depict estimates based on a closed population model with 2 visits to each plot. Gray ribbon indicates 95% credible intervals for estimates in each plot. Thick solid line is true number of nests in each plot (n = 5)

## Discussion

Our study is the first to apply mark-recapture methods to estimate the density of bumble bee nests, which represent a critical and understudied life cycle stage for this important pollinator group. In our study, single surveys of bumble bee nest abundance were subject to considerable bias and observation error, owing to imperfect detection. On average, we only detected 30% of existing nests on each 2-hour survey of a 0.3 ha plot, and we show that the low detection probability on single surveys can introduce substantial and spurious variation into counts (Fig. 4). Thus, in order to understand the nesting ecology and monitoring requirements of bumble bees, imperfect detection of nests must be properly accounted for.

Our estimate of detection probability is well within the range of reported rates for surveys of other sessile organisms: 0.01 to 1.0 in plants (Chen et al. 2013; Kellner and Swihart 2014), 0.17 to 0.60 for insect nests (Berberich et al. 2016; Brown et al. 2017), and 0.09 to 0.93 for patches of freshwater mussels (Reid 2016). These studies use mark-recapture approaches to improve estimates of density or occupancy on the landscape for sessile organisms that are not perfectly detectable, and our study adds bumble bees to this list of taxa.

The range of bumble bee nest densities we detected are comparable to those reported in Osborne et al. (2008), who used intensive fixed searches by citizen science volunteers to count nests in UK gardens and countryside habitats. Of our three plots, the highest density we detected was 33.3 nests·ha^−1^, similar to hedgerow (29.5 nests·ha^−1^), garden (35.8 nests·ha^−1^), and fence line (37.2 nests·ha^−1^) habitats reported in Osborne et al. (2008). Notably, Cumber (1953) is the only other study to report higher nest densities than these; his estimate of 48.6 nests·ha^−1^ was based on intensive free searches of a refuse dump in England. Conversely, our lowest density plot contained 10 nests·ha^−1^, which is similar to the density of 10.9 nests·ha^−1^ reported in Harder (1986) who intensively surveyed a successional field in Ontario, Canada. This estimate is also similar to the lowest densities in Osborne (10.8 nests·ha^−1^ in woodland and 11.4 nests·ha^−1^ in short grassland habitat). Therefore, our three plots seem to have captured the range of nest densities observed in other studies, if we restrict these studies to those with intensive search effort and extremely high detection probabilities.

Other studies have reported far lower nest densities than those in our study or those in Osborne et al. (2008), Cumber (1953), and Harder (1986). However, comparisons among studies are ultimately hampered by differences in survey efforts, and thus, differences in detection error. For example, low-intensity free searches by researchers or volunteers produced estimates of nest density ranging from 1.4 to 3.6 nests·ha^−1^ similar to the range of nest densities discovered by bumble bee “sniffer dogs” (O’Connor et al. 2012, 2017). Both of these studies acknowledge that detection error is likely substantial for these methods. Molecular studies also typically yield estimates of nest density that are 1-2 orders of magnitude lower than intensive ground-based searches (range: 0.13 to 1.9 nests·ha^−1^; Supplement 2). Several molecular studies have used ad-hoc approaches to account for imperfect detection, but these approaches likely under-estimate the true nest density (Goulson et al. 2010). Molecular methods also integrate habitat quality over larger spatial extents than ground-based surveys, and likely incorporate areas that are unsuitable for nesting (e.g., water bodies). Formal mark-recapture approaches are necessary to understand the degree to which variation in nest density between studies is driven by ecologically relevant factors (e.g., variation in habitat quality, differences in spatial scale at which studies occur) versus unresolved differences in imperfect detection.

In addition to estimating overall probability of nest detection, we found that nest detection declined when surveys were conducted later in the day and in warmer temperatures (Fig. 1e). Based on estimates from model *M*_*hour*_, mean detection probability during 6 am surveys was 0.40 (95% CRI = 0.31 to 0.50), but was only 0.05 (95% CRI = 0.01 to 0.17) for surveys initiated at 6 pm. This result is consistent with Kwon and Saeed (2003) who found that colony traffic and foraging activity of *Bombus terrestris* declined throughout the day and when temperatures were warmer. Similarly, Couvillon et al. (2010) reported that workers of all sizes conducted fewer foraging trips in warmer temperatures. Thus, the results of our mark-recapture estimate of detection probability are broadly consistent with prior knowledge of bumble bee foraging ecology, and suggest that variation in forager behavior is a likely driver of differences in detectability.

Although we did not measure them in our study, other factors could also influence colony activity, and in turn, the probability that nests are detected on a given survey. For example, larger colonies have higher traffic at nest entrances (Kwon and Saeed 2003) and are therefore likely to be more detectable. Colony size, in turn, depends on floral resources available throughout the season (Williams et al. 2012; Crone and Williams 2016). Thus, detection of nests may differ between high and low quality habitat owing to systematic differences in colony size. Here, we found evidence for systematic differences in nest detection across our three survey plots. Plot 1 had the highest nest density (33.3 nests·ha^−1^), and simultaneously, detection was highest for nests in this plot (Fig. 1d). Mark-recapture approaches can correct for these spatio-temporal biases in detection probability, which in turn, improve comparisons of abundance within and between studies.

Population monitoring schemes will often have multiple objectives, in addition to generating reliable estimates of population size (e.g., sampling large numbers of individuals to assess body condition, biochemistry, or disease status; monitoring behavior of individual animals in different environmental contexts). For rare or highly cryptic species, intensive surveys over a small area may be required to generate reliable estimates of local abundance, but yield few individuals for further detailed study. In the case of bumble bees, intensive fixed searches of small plots are considered sufficient to detect all existing nests and therefore reliably estimate nest density (Osborne *et al.*, 2008; O’Connor et al., 2012). However, intensive fixed searches are inefficient when a large number of nests are desired (e.g., for examining microsite characteristics associated with nests, or for studying workers at nest entrances) and are logistically challenging to implement over large areas without distributed citizen science networks (Osborne et al. 2008; Lye et al. 2012). We were able to locate a high number of nests, generate precise estimates of abundance with only 5-6 surveys in each of our 3 plots (Fig. 3a), and produce relatively unbiased estimates with an even smaller number of surveys at a larger number of sites (Fig 4). This equates to much lower search effort than typical intensive fixed searches, while simultaneously locating nests at a comparable rate to low-intensity free searches (O’Connor et al. 2012). Our study illustrates the advantage of mark-recapture for optimizing survey protocols for cryptic and sessile organisms. Further research in this area will be valuable in illuminating the ecological drivers of pollinator nesting ecology, a critical but understudied subject.

## Authors’ Contributions

All authors conceived of research ideas, designed methodology, and collected field data. DI and EC led statistical analysis and writing of the manuscript. All authors contributed critically to the drafts and gave final approval for publication.

## Data Accessibility

All data and code associated with these analyses will be archived in Dryad Digital Repository upon acceptance of this manuscript.

## Compliance with Ethical Standards

The authors declare that they have complied with ethical standards.

## Acknowledgements

We thank the Trustees of Reservations and Appleton Farms for providing access to their properties on which this study was conducted. We thank Russ Hopping for invaluable assistance with study design. We also thank Annika Greenleaf, Max McCarthy, Moshe Steyn, Erin Wampole, and our sniffer-dogs-in-training, Indy and Molly, for assistance with field work. This work was supported by the US National Science Foundation (DEB1354022) and the US Strategic Environmental Research and Development Program (SERDP, RC-2119).

## Appendix 1

**Table 2.**
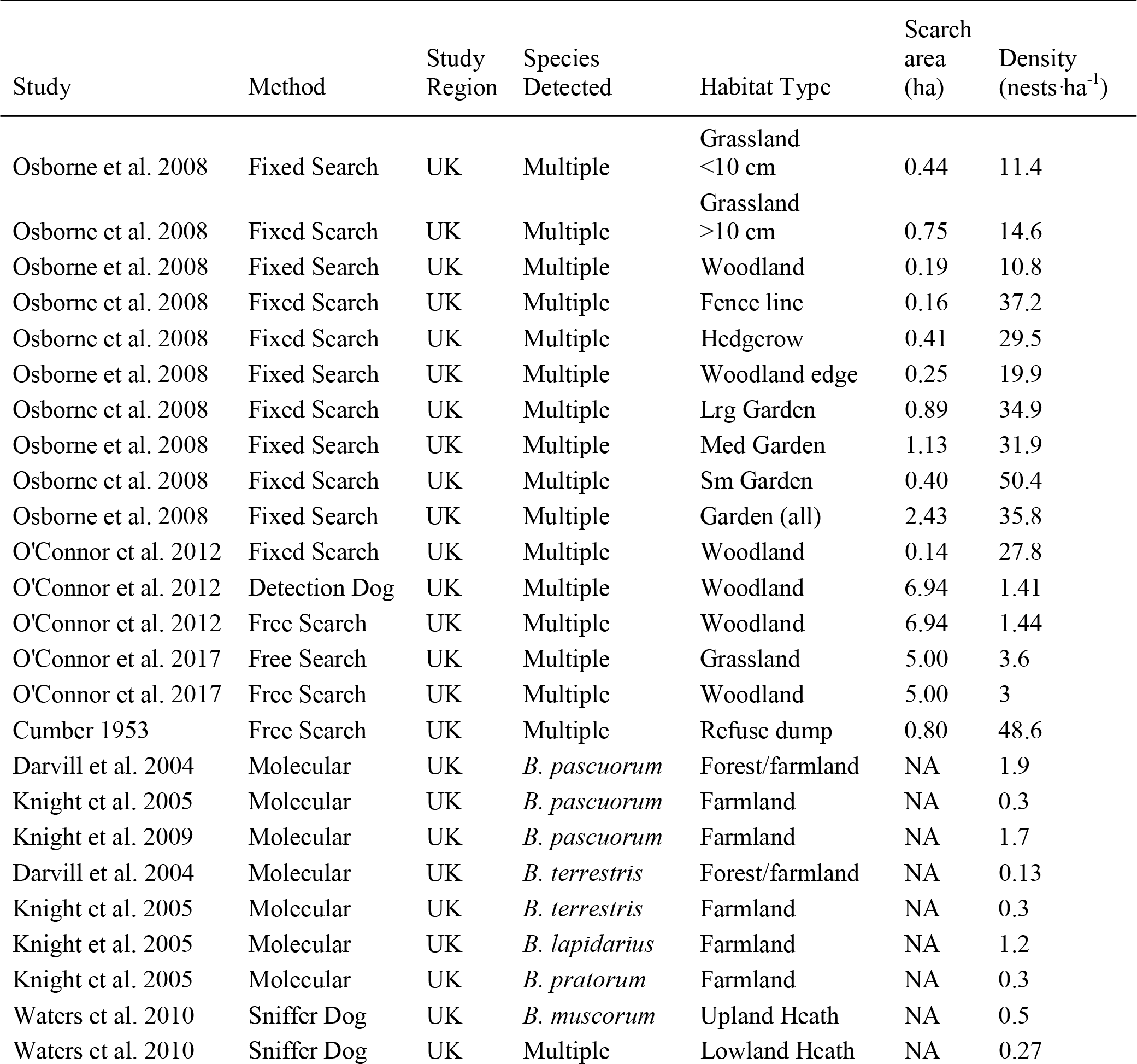
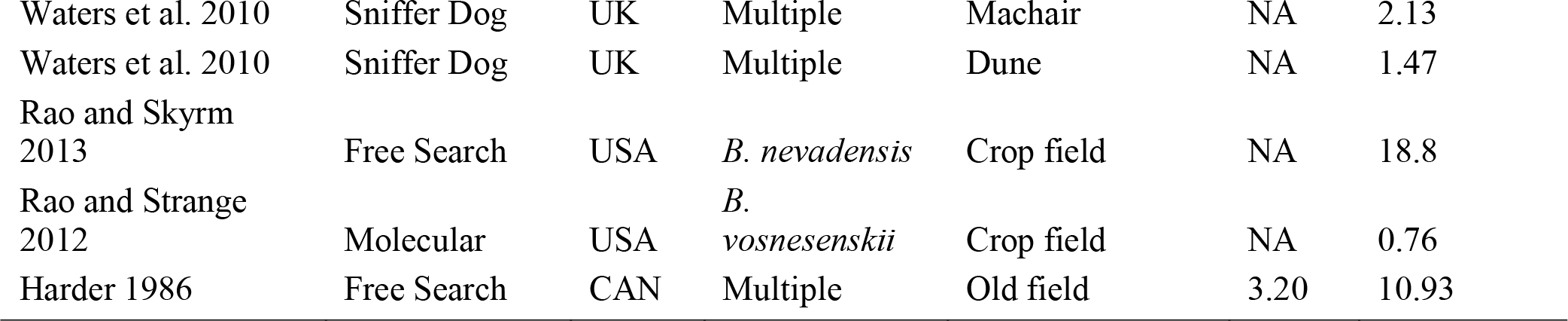
Summary of previous studies that estimated bumble bee nest density, with description of methods used, study region, species detected, habitat types associated with density estimates, area searched, and estimated density. Note that search area does not apply to molecular methods as nest density is inferred from genetic relatedness between workers and estimates of worker foraging range

## Description of supplementary information

Supplementary material 1: R code used to examine variation in the probability of detecting bumble bee nests using bayesian closed population models and generate Fig. 1 and Fig. 2

Supplementary material 2: R code used to demonstrate that imperfect detection can obscure differences in colony density among plots and generate Fig. 3

Supplementary material 3: R code used to estimate bumblebee populations for simulated mark-recapture data and generate Fig. 4

Supplementary material 4: Capture histories for 10 bumble bee nests for survey plot 1

Supplementary material 5: Capture histories for 5 bumble bee nests for survey plot 2

Supplementary material 6: Capture histories for 3 bumble bee nests for survey plot 3

Supplementary material 7: Covariate data for each nest, including the height of the tallest vegetation immediately above the nest entrance

Supplementary material 8: Hourly ambient air temperature measurements during each survey from the nearest weather station

## References

Anderson DR (2001) The need to get the basics right in wildlife field studies. Wildl Soc Bull (1973-2006) 29:1294–1297

Berberich GM, Dormann CF, Klimetzek D, Berberich MB, Sanders NJ, Ellison AM (2016) Detection probabilities for sessile organisms. Ecosphere 7:e01546. https://doi.org/10.1002/ecs2.1546

Brown LM, Breed GA, Severns PM, Crone EE (2017) Losing a battle but winning the war: moving past preference–performance to understand native herbivore–novel host plant interactions. Oecologia 183:441–453. https://doi.org/10.1007/s00442-016-3787-y

Chen G, Kéry M, Plattner M, Ma K, Gardner B (2013) Imperfect detection is the rule rather than the exception in plant distribution studies. J Ecol 101:183–191. https://doi.org/10.1111/1365-2745.12021

Couvillon MJ, Fitzpatrick G, Dornhaus A (2010) Ambient air temperature does not predict whether small or large workers forage in bumble bees (Bombus impatiens). Psyche: A Journal of Entomology 2010:536430. http://dx.doi.org/10.1155/2010/536430

Crone EE, Williams NM (2016) Bumble bee colony dynamics: quantifying the importance of land use and floral resources for colony growth and queen production. Ecol Lett 19:460–468. http://dx.doi.org/10.1111/ele.12581

Cumber R (1953) Some aspects of the biology and ecology of humble-bees bearing upon the yields of red-clover seed in New Zealand. New Zealand Journal of Science and Technology 34:227–240

Darvill B, Knight ME, Goulson D (2004) Use of genetic markers to quantify bumblebee foraging range and nest density. Oikos 107:471–478. https://doi.org/10.1111/j.00301299.2004.13510.x

Goulson D (2010) Bumblebees: behaviour, ecology, and conservation, 2nd edn. Oxford University Press, New York, pp 317

Goulson D, Lepais O, O’Connor S, Osborne JL, Sanderson RA, Cussans J, Goffe L, Darvill B (2010) Effects of land use at a landscape scale on bumblebee nest density and survival. J Appl Ecol 47:1207–1215. https://doi.org/10.1111/j.1365-2664.2010.01872.x

Gu W, Swihart RK (2004) Absent or undetected? Effects of non-detection of species occurrence on wildlife–habitat models. Biol Conserv 116: 195–203. https://doi.org/10.1016/S0006-3207(03)00190-3

Harder LD (1986) Influences on the density and dispersion of bumble bee nests (Hymenoptera: Apidae). Ecography 9:99–103. https://doi.org/10.1111/j.1600-0587.1986.tb01196.x

Heard M, Carvell C, Carreck N, Rothery P, Osborne J, Bourke A (2007) Landscape context not patch size determines bumble-bee density on flower mixtures sown for agri-environment schemes. Biol Lett 3:638–641. https://doi.org/10.1098/rsbl.2007.0425

Herrmann F, Westphal C, Moritz RF, Steffan□ Dewenter I (2007) Genetic diversity and mass resources promote colony size and forager densities of a social bee (Bombus pascuorum) in agricultural landscapes. Mol Ecol 16:1167–1178. https://doi.org/10.1111/j.1365294X.2007.03226.x

Kellner KF, Swihart RK (2014) Accounting for imperfect detection in ecology: a quantitative review. PLoS ONE 9:e111436. https://doi.org/10.1371/journal.pone.0111436

Kellner KF (2018) jagsUI: A Wrapper Around ‘rjags’ to Streamline ‘JAGS’ Analyses. R package version 1.5.0. https://CRAN.R-project.org/package=jagsUI

Kells AR, Goulson D (2003) Preferred nesting sites of bumblebee queens (Hymenoptera: Apidae) in agroecosystems in the UK. Biol Conserv 109:165–174. https://doi.org/10.1016/S0006-3207(02)00131-3

Kéry M, Schaub M (2012) Bayesian population analysis using WinBUGS: a hierarchical perspective. Academic Press, Cambridge

Kéry M, Schmidt B (2008) Imperfect detection and its consequences for monitoring for conservation. Community Ecol 9:207–216. https://doi.org/10.1556/ComEc.9.2008.2.10

Kwon YJ, Saeed S (2003) Effect of temperature on the foraging activity of Bombus terrestris L.(Hymenoptera: Apidae) on greenhouse hot pepper (Capsicum annuum L.). Appl Entomol Zool 38:275–280. https://doi.org/10.1303/aez.2003.275

Lye GC, Osborne JL, Park KJ, Goulson D (2012) Using citizen science to monitor Bombus populations in the UK: nesting ecology and relative abundance in the urban environment. J Insect Conserv 16:697–707. https://doi.org/10.1007/s10841-011-9450-3

O’Connor S, Park KJ, Goulson D (2012) Humans versus dogs; a comparison of methods for the detection of bumble bee nests. J Apicult Res 51:204–211. https://doi.org/10.3896/IBRA.1.51.2.09

O’Connor S, Park KJ, Goulson D (2017) Location of bumblebee nests is predicted by counts of nest-searching queens. Ecol Entomol 42:731–736. https://doi.org/10.1111/een.12440

Osborne JL, Martin AP, Shortall CR, Todd AD, Goulson D, Knight ME, Hale RJ, Sanderson RA (2008) Quantifying and comparing bumblebee nest densities in gardens and countryside habitats. J Appl Ecol 45:784–792. https://doi.org/10.1111/j.1365-2664.2007.01359.x

Otis DL, Burnham KP, White GC, Anderson DR (1978) Statistical inference from capture data on closed animal populations. Wildl Monogr 62: 3–135

Rao S, Skyrm KM (2013) Nest Density of the Native Bumble Bee, Bombus nevadensis Cresson (Hymenoptera: Apoidea), in an Agricultural Landscape. J Kans Entomol Soc 86:93–97. https://doi.org/10.2317/JKES120708.1

Reid S (2016) Search effort and imperfect detection: Influence on timed-search mussel (Bivalvia: Unionidae) surveys in Canadian rivers. Knowl Manag Aquat Ecosyst 1–8. https://doi.org/10.1051/kmae/2016004

Slade NA, Alexander HM, Dean Kettle W (2003) Estimation of population size and probabilities of survival and detection in Mead’s milkweed. Ecology 84:791–797. https://doi.org/10.1890/0012-9658(2003)084[0791:EOPSAP]2.0.CO;2

Svensson B, Lagerlöf J, Svensson BG (2000) Habitat preferences of nest-seeking bumble bees (Hymenoptera: Apidae) in an agricultural landscape. Agr Ecosyst Environ 77:247–255. https://doi.org/10.1016/S0167-8809(99)00106-1

Waters J, O’Connor S, Park KJ, Goulson D (2010) Testing a detection dog to locate bumblebee colonies and estimate nest density. Apidologie 42:200–205. https://doi.org/10.1051/apido/2010056

White GC, Burnham KP (1999) Program MARK: survival estimation from populations of marked animals. Bird Study 46:S120–S139. https://doi.org/10.1080/00063659909477239

Williams BK, Nichols JD, Conroy MJ (2002) Analysis and management of animal populations. Academic Press

Williams NM, Regetz J, Kremen C (2012) Landscape□scale resources promote colony growth but not reproductive performance of bumble bees. Ecology 93:1049–1058. https://doi.org/10.1890/11-1006.1

