## Supplementary material for "Accounting for imperfect detection in species with sessile life cycle stages: a case study of bumble bee colonies": ESM_9.txt

        model{

        ########
        # Priors
        ########
        mean.p ~ dunif(0, 1) # Capture/recapture probability on 0 to 1 scale
        mean.lp <- log(mean.p / (1-mean.p)) # logit scale

        omega ~ dunif(0, 1) # Probability of true presence of each individual in augmented dataset

        ########
        # Likelihood
        ########
        for (i in 1:M){

        # True presence of individuals
        z[i] ~ dbern(omega)

        lp[i] <- mean.lp
        p[i]  <- 1 / (1 + exp(-lp[i]))  # Inverse logit (place back on 0 to 1 scale)

        for (j in 1:T){

        p.eff[i,j] <- z[i] * p[i] # Can only be detected if z = 1 (truly present); if z = 0 (truly absent) then detection must be 0

        y[i,j] ~ dbern(p.eff[i,j]) # Was individual detected or not?

        } #j
        } #i

        ########
        # Derived quantities
        ########
        N <- sum(z[]) # Sum up the number of *truly present* individuals to give estimate of abundance
        }
        
