## Supplementary material for "Accounting for imperfect detection in species with sessile life cycle stages: a case study of bumble bee colonies": Metadata.docx

ESM_1.txt

Supplementary material 1: R code used to examine variation in the probability of detecting bumble bee nests using bayesian closed population models

ESM_2.txt

Supplementary material 2: R code used to demonstrate that imperfect detection can obscure differences in colony density among plots

ESM_3.txt

Supplementary material 3: R code used to estimate bumblebee populations for simulated mark-recapture data

ESM_4.csv

Supplementary material 4: Capture histories for 10 bumble bee nests for survey plot 1

ESM_5.csv

Supplementary material 5: Capture histories for 5 bumble bee nests for survey plot 2

ESM_6.csv

Supplementary material 6: Capture histories for 3 bumble bee nests for survey plot 3

ESM_7.csv

Supplementary material 7: Covariate data for each nest, including the height of the tallest vegetation immediately above the nest entrance

ESM_8.csv

Supplementary material 8: Hourly ambient air temperature measurements during each survey from the nearest weather station

ESM_9.txt

Supplementary material 9: JAGS script to run simulations (ESM_3)
